# TKGWV2: An ancient DNA relatedness pipeline for ultra-low coverage whole genome shotgun data

**DOI:** 10.1101/2021.06.22.449449

**Authors:** Daniel M. Fernandes, Olivia Cheronet, Pere Gelabert, Ron Pinhasi

## Abstract

Estimation of genetically related individuals is playing an increasingly important role in the ancient DNA field. In recent years, the numbers of sequenced individuals from single sites have been increasing, reflecting a growing interest in understanding the familial and social organisation of ancient populations. Although a few different methods have been specifically developed for ancient DNA, namely to tackle issues such as low-coverage homozygous data, they require a 0.1 - 1x minimum average genomic coverage per analysed pair of individuals between. Here we present an updated version of a method that enables estimates of 1st and 2nd-degrees of relatedness with as little as 0.026x average coverage, or around 1.3 million aligned reads per sample - 4 times less data than 0.1x. By using simulated data to estimate false positive error rates, we further show that a threshold even as low as 0.012x, or around 600,000 reads, will always show 1st-degree relationships as related. Lastly, by applying this method to published data, we are able to identify previously undocumented relationships using individuals previously excluded from kinship analysis due to their very low coverage. This methodological improvement has the potential to enable relatedness estimation on ancient whole genome shotgun data during routine low-coverage screening, and therefore improve project management when decisions need to be made on which individuals are to be further sequenced.

## Introduction

The estimation of genetic relatives in ancient DNA (aDNA) research has become an integral part of any studies that involve individuals from the same site or region. The challenging nature of aDNA has led researchers in recent years to adapt traditional relatedness estimation methods, such as those present in PLINK^1^ and KING^2^, to datasets of ancient individuals for which diploid and high-quality data is not available. This has resulted in the development of a few different methods, using pseudo-haploid data and/or genotype likelihoods, that have been shown to work with genomic coverages as low as between 0.1 and 1x^3–7^.

Inferring these relationships is essential, not only for addressing questions related to social and familial organisation of ancient groups, but also as a quality control step for population-based analyses, where an excess of very close relatives may introduce biases that reflect higher rates of allele sharing among individuals than would be expected among non-relatives, and, therefore, do not represent the real genetic variation of the whole population^8^. Specifically, one individual from a pair of 1st-degree relatives is routinely excluded from analyses requiring the grouping of individuals to avoid these biases^9^, which leads to a potential very limited use of otherwise good data from the excluded individual.

With this in mind, individual aDNA projects would be able to better structure their workflow and budget attributions if kinship relationships could be estimated at early stages of the research plan -such as during ultra-low-coverage screening, which is a cheap and effective way to evaluate the quantity and quality of data that can be expected from an ancient individual. For projects where the analysis of kinship and related individuals is the main focus, researchers would be able to concentrate their resources on particular individuals targeted for the specific research questions. As aDNA laboratories routinely screen their samples using ultra-low-coverage sequencing data before deciding which samples or individuals are to be further sequenced into the desired depths, having information on kinship during this phase could be an advantage.

However, the most widely used relatedness estimation methods/software in aDNA research typically require substantially larger amounts of data than that which originates in routine screening runs, which typically yield only low amounts of data. For example, a 0.1x genomic coverage threshold has been used in the past in software such as NgsRelate^6^ and READ^3^, in order to confidently estimate 1st and 2nd-degree relationships. Another commonly used software, lcMLkin, in turn requires between and 1x coverage to identify 1st and 2nd-degree relatives, or at least 10,000-20,000 common single nucleotide polymorphisms (SNPs), after quality control^5^.

Here, we present TKGWV2 (“Thomas Kent Genome-Wide Variants 2”), an update to a method published in 2017^4^ that, by using genome-wide variants instead of variant sets commonly used in aDNA research, such as the 1240K Capture or Affymetrix Human Origins arrays, increases the amount of potentially available data for the method’s relatedness estimator from 1,240,000 or 600,000 to over 22,000,000 non-fixed biallelic variants present in the 1000 Genomes Project Phase 3^10^ -a 18 to 37 times gain, respectively. As a result, the amount of whole genome shotgun sequencing data required to estimate at least 1st and 2nd-degree relationships is reduced by at least ∼4 times, potentially allowing the estimation of close relatedness between ancient individuals during routine screening stages.

## Results

### Method’s pipeline description

TKGWV2 can be publicly accessed at https://github.com/danimfernandes/tkgwv2. It requires the user to provide three types of files: aligned individual *.BAM files, a list of non-fixed biallelic SNPs for genotype calling, and the population allele frequencies of those same SNPs. These are then processed in three consecutive steps:

1.Genotype calling and conversion of pileup files into individual pseudo-haploid PLINK text files;

2.Identification of overlapping variants per pairs of individuals, extraction of corresponding allele frequencies from the provided frequencies file, and creation of transposed PLINK text files for the next step;

3.Pairwise relatedness estimation (Rxy).

We use Queller & Goodnight’s^11^ estimator and therefore require population-wide allele frequencies for the SNP set used. If the input data is already in a genotype dataset format such as PLINK, TKGWV2 can be started from point 2 and will only require the allele frequencies file. The only quality control step required for the selection of SNPs is to include only non-fixed biallelic variants. Low frequency variants do not need to be excluded, meaning that an overall higher number of SNPs is available for the estimator. Three relatedness classes are evaluated: 1st-degree, 2nd-degree, and unrelated. However, due to the use of pseudo-haploid data, as commonly seen in very low coverage ancient shotgun data, the resulting relatedness coefficients (r) are half of the original expected values, and therefore we refer to them as “halved relatedness coefficients” (HRC). The three equidistant relatedness classes used in our method have an average HRC of 0.25 for 1st-degree relatives, 0.125 for 2nd-degree relatives, and 0 for unrelated individuals, and following their normal distribution ranges in real populations, we set hard thresholds for each class’s range of values at the mid-points between classes -0.1875 between 1st-and 2nd-degree relationships, and 0.0625 for the latter and unrelated.

### Application to known relatives from the 1000 Genomes Phase 1

To validate this new approach to the method presented in Fernandes et al.^4^, we used the publicly available data from the 1000 Genomes Phase 1, which includes individuals with known relationships^10^. The downloadable VCF data for ∼38 million SNPs and indels was converted into PLINK format, and only non-fixed biallelic SNPs were used during analysis. We chose the Southern Han Chinese (CHS) population as it includes 10 known 1st-or 2nd-degree relationships shared between 13 individuals. All remaining 4,940 pairwise relationships are reported as unrelated. We started by subsampling the 100 CHS individuals at 8 fractions between 0.5% and 10% of the complete 38 million variant coverage. We did not specify a minimum number of used SNPs in order to test the total variation of the estimates.

In the fraction corresponding to a subsampling of 2%, which averaged 4,059 SNPs used per test, all 10 known relationships described in Phase 1 of the 1000 Genomes Project were correctly identified and assigned the correct degree of relatedness (**Figure 1**). However, in this fraction a few unrelated individuals also crossed the threshold for 2nd-degree of relatedness at 0.0625. These potential false positives seem to be corrected from the next fraction onwards (3% subsampling fraction, average 9,137 SNPs), and so all expected relationships are correctly assigned. Only the pair of individuals HG00475 and HG00542 consistently shows an HRC within the 2nd-degree class, and although they were not detected or declared as relatives in the 1000 Genomes project, previously published research has already described them as 2nd-degree relatives^12^. These results suggest that, for the CHS population, the error rates are likely to be very low starting from around 10,000 shared SNPs.

**Figure 1:**
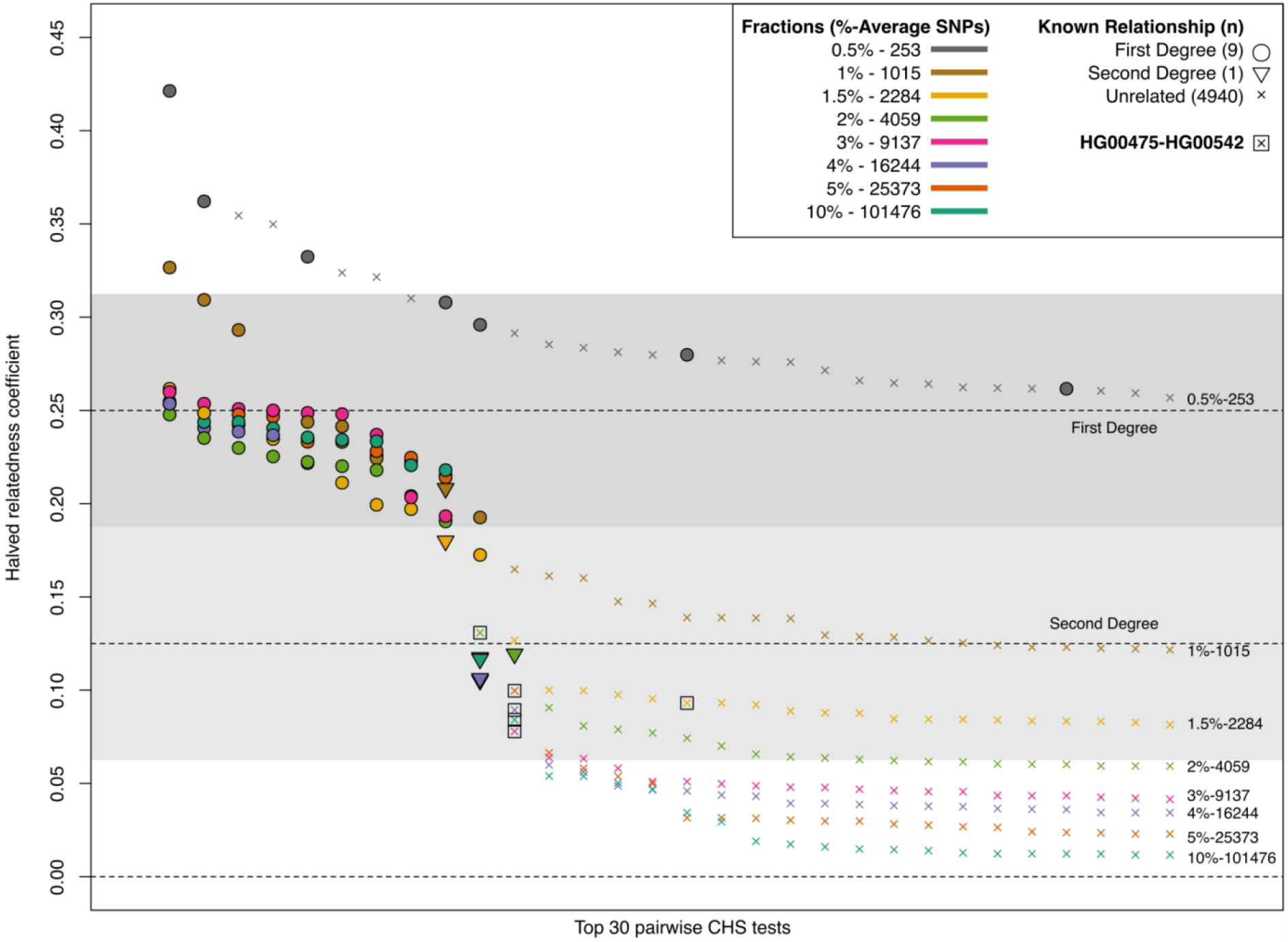
Top 30 results ordered by HRC of all 4,950 pairwise tests for the 100 individuals from the CHS population, ordered by relatedness coefficient, for the 8 subsampling fractions between 0.5 and 10%. Known relationships are shown as filled symbols, and each different fraction as a specific colour. All triangles (second degree) are expected to fall within the lighter gray area, and all circles (first degree) within the darker gray area. The allele frequencies from the CHS population were used.

### Application to published ancient DNA data

We next applied this approach to an ancient shotgun dataset from Schroeder et al.^13^ (ENA accession number PRJEB28451), composed of 15 Late Neolithic individuals from a Globular Amphora mass grave from the site of Koszyce, in present-day Poland. The unusually high number of genetic relatives of up to 3rd-degree detected at this site (n=55) make it a great candidate for testing our new pipeline. However, since 3rd-degree relationships (0.0625 average HRC) are expected to be represented by major overlaps with the unrelated and 2nd-degree relative classes, TKGWV2 only assesses up to 2nd-degree relationships. For this reason, we did not consider the 11 pairs of 3rd-degree relatives identified by Schroeder et al.^13^ as part of our pass/fail assessments. We used 1000 Genomes Phase 3 modern day allele frequencies from 503 individuals with European ancestry^10^, restricted to ∼22 million non-fixed biallelic SNPs.

This time, instead of randomly subsampling each individual’s genotypes, we investigated genomic coverage thresholds for the application of this method to shotgun sequenced individuals by subsampling the BAM files of the 15 Koszyce individuals to specific numbers of aligned and non-duplicated reads: 3,000,000, 2,500,000, 2,000,000, 1,600,000, 1,300,000, 1,000,000, 800,000, 600,000, 400,000, and 200,000 (**Supplementary Table 1)**. At 1,300,000 aligned reads per sample (corresponding to about 0.026x genomic coverage and an average of 18,364 SNPs shared per pairwise test), all unrelated individuals were assigned to the correct relatedness class (HRC below 0.0625), only two out of 56 2nd-degree relative pairs were incorrectly assigned with a HRC below 0.0625 (individuals 11-9 and 14-4), and only one pair of 1st-degree relative pairs (individuals 10-11) has a HRC under the 0.1875 threshold for 1st-degree relatives (0.1553) (**Figure 2, Supplementary Table 2**). In the next higher subset, at 1,600,000 aligned reads (about 0.032x genomic coverage and an average of 27,589 SNPs shared), again all unrelated individuals were correctly assigned, while two 2nd-degree relative pairs had a HRC out of their range (individuals 11 and 9 with 0.0547 and individuals 10 and 15 with 0.1909). All 1st-degree relative pairs were correctly identified and had a HRC above 0.1875 (**Figure 2, Supplementary Table 2**).

**Table 1:**
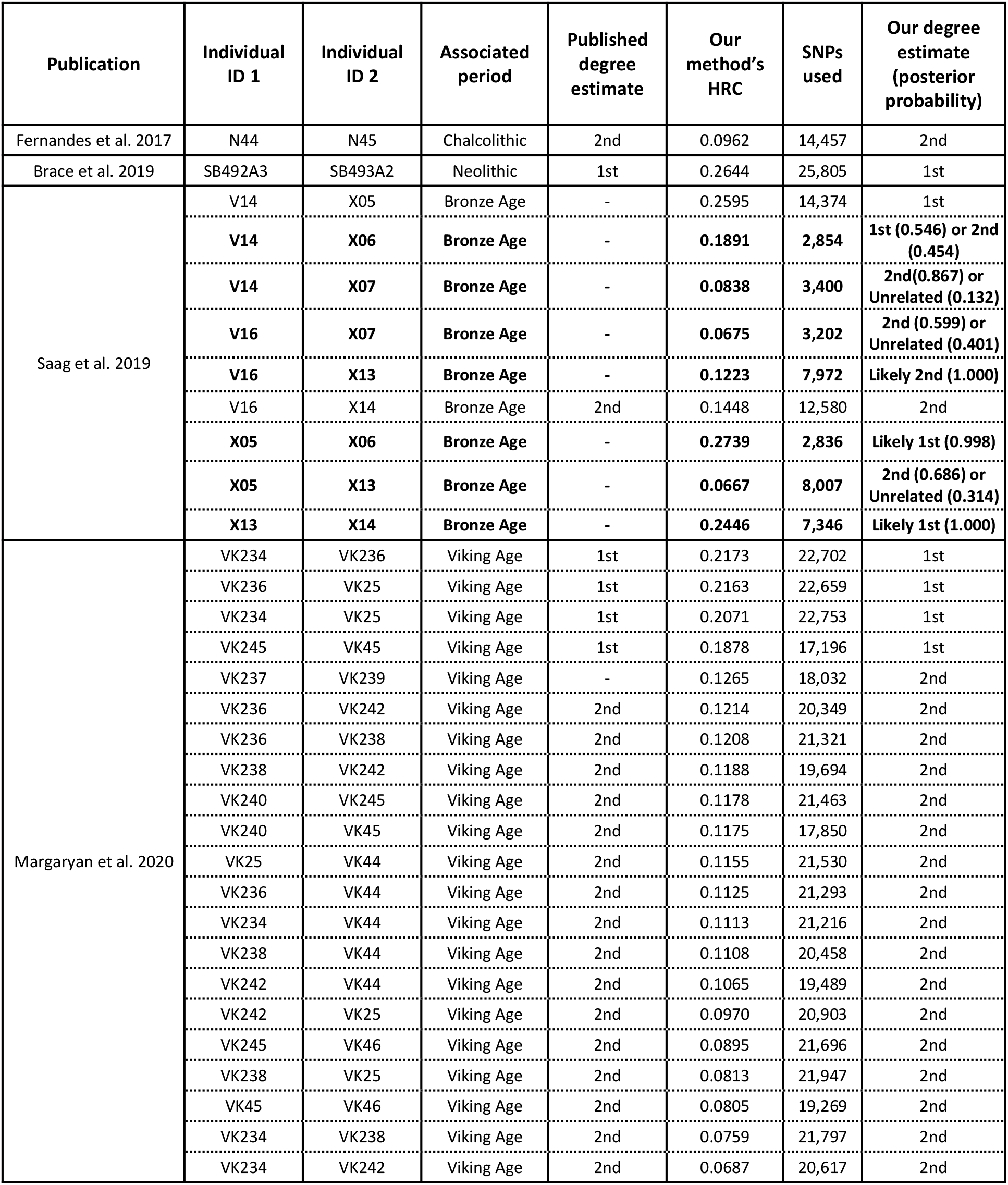
Application of the method to other published ancient individuals. BAM files were downloaded from the European Nucleotide Database, subsampled to a maximum of 1,300,000 reads, and then processed through our pipeline. The allele frequencies used were from individuals with European ancestry in the 1000 Genomes Phase 3 dataset. Estimates in bold are based on less than 10,000 SNPs, and therefore include some degree of uncertainty. For these, we present the posterior probabilities of each degree between parentheses, and on **Supplementary Figure 2** we show the corresponding simulated range plots for these pairs.

**Figure 2:**
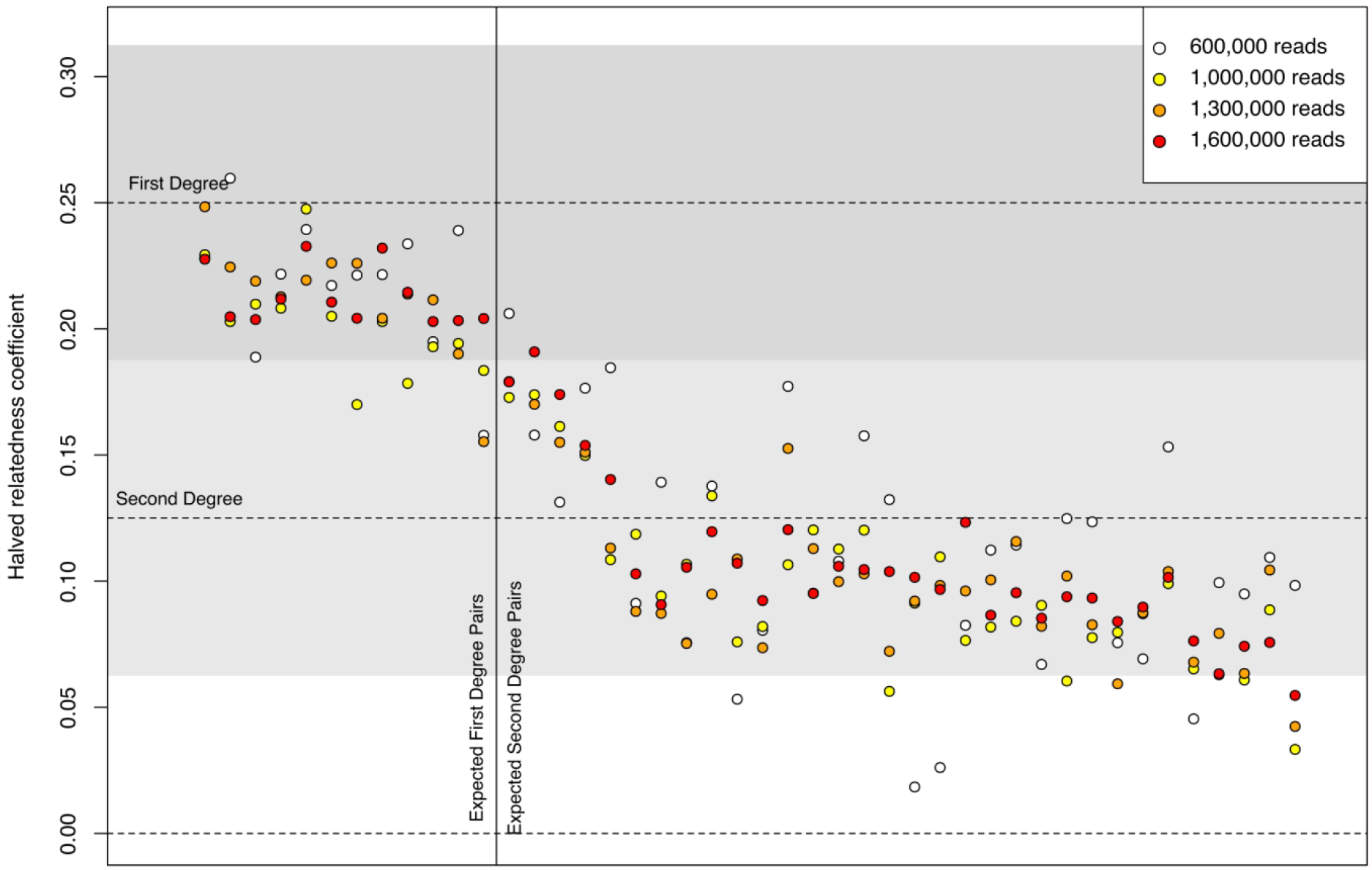
Coefficients of the 44 relationships for 1st-and 2nd-degree relatives from the Neolithic site of Koszyce^13^. Each vertical set of 4 coloured points represents one relationship tested with different numbers of aligned reads, according to the legend. Relationships to the left of the solid horizontal line are expected to be 1st-degree, as per Schroeder et al.^13^, and relationships to the right of the line are expected to be 2nd-degree. Grey areas define 1st-and 2nd-degree range intervals.

These results suggest that from around 18,000 SNPs (1,300,000 reads, or 0.026x coverage) the rate of incorrectly assigned relationships is ∼3.2%, at 3 out of 94 pairwise tests. However, this number was consistent even up to around 93,000 SNPs and almost always involved individuals 5, 10, 11, and 15, whose pairwise HRCs were always in the region between 1st-and 2nd-degrees. In their original publication, these individuals were identified as possible half-siblings who shared a biological father, and possibly different mothers that could be related to each other to varying degrees, due to sharing the same mitochondrial haplogroup. If this is indeed the case, these parental relationships would be consistent with producing individuals with an intermediate coefficient between 1st-and 2nd-degrees (¾-siblings, 0.1875 HRC), which would be in agreement with our results.

Since TKGWV2 does not assess 3rd-degree relationships, these will show up as either 2nd-degree or unrelated. On **Supplementary Table 2** we show that from 18,000 SNPs used, the 11 3rd-degree relationships identified by Schroeder and colleagues^13^ range between -0.0095 and 0.0931 HRC (0.0517 average over 55 tests). Conservatively, then, 2nd-degree relationships between 0.0625 and

∼0.090 may instead be referred to as “2nd-or 3rd-degree” to cover this possibility.

Although we argue that 1,300,000 reads (or 0.026x coverage) could be used as a minimum threshold for the application of this method with very low error rates, we also noticed that from around 1,000,000 reads (11,000 SNPs or 0.020x coverage) all true unrelated individuals had an HRC below 0.0625, indicating that this lower threshold can possibly be used to avoid false positive related pairs, as any pair above 0.0625 would necessarily share a 1st-or 2nd-degree relationship. This would agree with the CHS results shown in **Figure 1**, where at the 3 and 4% fractions (9,137 and 16,244 SNPs) all unrelated individuals also began to have a HRC below 0.0625. On the other hand, false negatives at this 0.020x coverage are possible, as at least four 2nd-degree pairs from Koszyce had a HRC below 0.0625 (**Supplementary Table 2**).

Lastly, by looking at even lower coverages, we noticed that true 1st-degree relatives showed as either 1st-or 2nd-degree from around 4,000 SNPs (600,000 reads or 0.012x coverage), with only a single pair of individuals below 0.1875 (individuals 10 and 11 -0.1578 HRC) (**Figure 2**; **Supplementary Table 2**).

To extend the method to individuals from different time periods, and therefore different ancestries and allele frequencies, we also applied TKGWV2 at the conservative 1,300,000 reads threshold to other published shotgun sequenced relatives, identifying the published degrees of relatedness with a 100% success rate (**Table 1**). However, to demonstrate the application of our pipeline to ultra low coverages, we also included some individuals from these publications that were not included in their original kinship analysis because of coverages below the defined thresholds. For the Church burial of the Faroe Islands in Margaryan et al.^14^ we included individuals VK239 and VK248 in the analysis, with coverages below the publication’s threshold of 0.1x (0.027x and 0.076x, respectively), and identified a new 2nd-degree relationship for VK239-VK237 (**Table 1**). In the case of Saag et al.^15^, for Toomani (Muuksi) we included individuals X05, X06, and X07, who have coverages below their threshold of 0.03x (0.0290x, 0.0048x, and 0.0059x, respectively). From Lastekangur (Rebala) we included X13 (0.0147x). Overall, we were able to confidently identify a new pair of 1st-degree relatives in Toomani (V14-X05, 0.2595 HRC, 14,374 SNPs), although 7 other relationships were also possibly identified (**Table 1**; in **Supplementary Fig. 1** we show a putative family tree for X13, X14, and V16, using this new data). Due to the lower coverage of the individuals involved in these relationships, the number of used SNPs ranged from 2,854 to 8,007, and therefore the degree of uncertainty and error rates are higher. We therefore used simulated data based on the allele frequencies of the population to attach posterior probabilities to each estimate -a process we describe in the section below.

### Simulated distribution ranges and class overlaps

In the tests above we used hard thresholds as a simplified way to bin the different coefficients into each class, and for the CHS population we saw that from around 10,000 SNPs all expected 1st-degree relatives, 2nd-degree relatives, and unrelated relationships were estimated correctly, whereas for the ancient population from around 18,000 SNPs a maximum of three relationships were incorrectly assigned. However, since the number of known relationships in any population is small, including within the 1000 Genomes Project, simulations are useful to better quantify false positive rates from distribution ranges and class overlaps, and to calculate posterior probabilities for each HRC. Moreover, although 1st-and 2nd-degree relatives are theoretically expected to share 50% and 25% of their variation (or 0.25 and 0.125 for HRC), respectively, the observed and estimated coefficients will be influenced by the reference population panel used and the background relatedness between all pairs of individuals within it^16^. Therefore, simulated distribution ranges are useful to represent this variability within the population.

We first generated 500 pairs of individuals of the unrelated, 1st-degree, and 2nd-degree classes, for an increasing range of randomly subset SNPs (**Figure 3**). For the subsets with the lower numbers of SNPs used (for example, 1,000 and 5,000), the overlap between the three HRC equidistant classes’ ranges is substantial, however, from between 10,000 and 20,000 SNPs the normal distributions tend towards having no overlaps (**Figure 3a**). This lack of overlaps is caused by the larger SNP datasets including more variability that is only captured when larger population sizes are simulated. Considering this, by increasing the number of simulated individuals from 500 to 5,000 we induce an expansion of the normal curves, and again show overlapping ranges between all classes (**Figure 3b**). As posterior probabilities cannot be assigned to relatedness estimates that fall on areas not covered by any curve, having overlapping ranges is essential, and as seen, the more SNPs used, the larger the number of simulated pairs required to fully cover this range. Assigning posterior probabilities to each class is especially useful when using sub-optimal SNP counts below the thresholds mentioned previously, as the resulting estimates will have higher uncertainty and larger overlaps.

**Figure 3.**
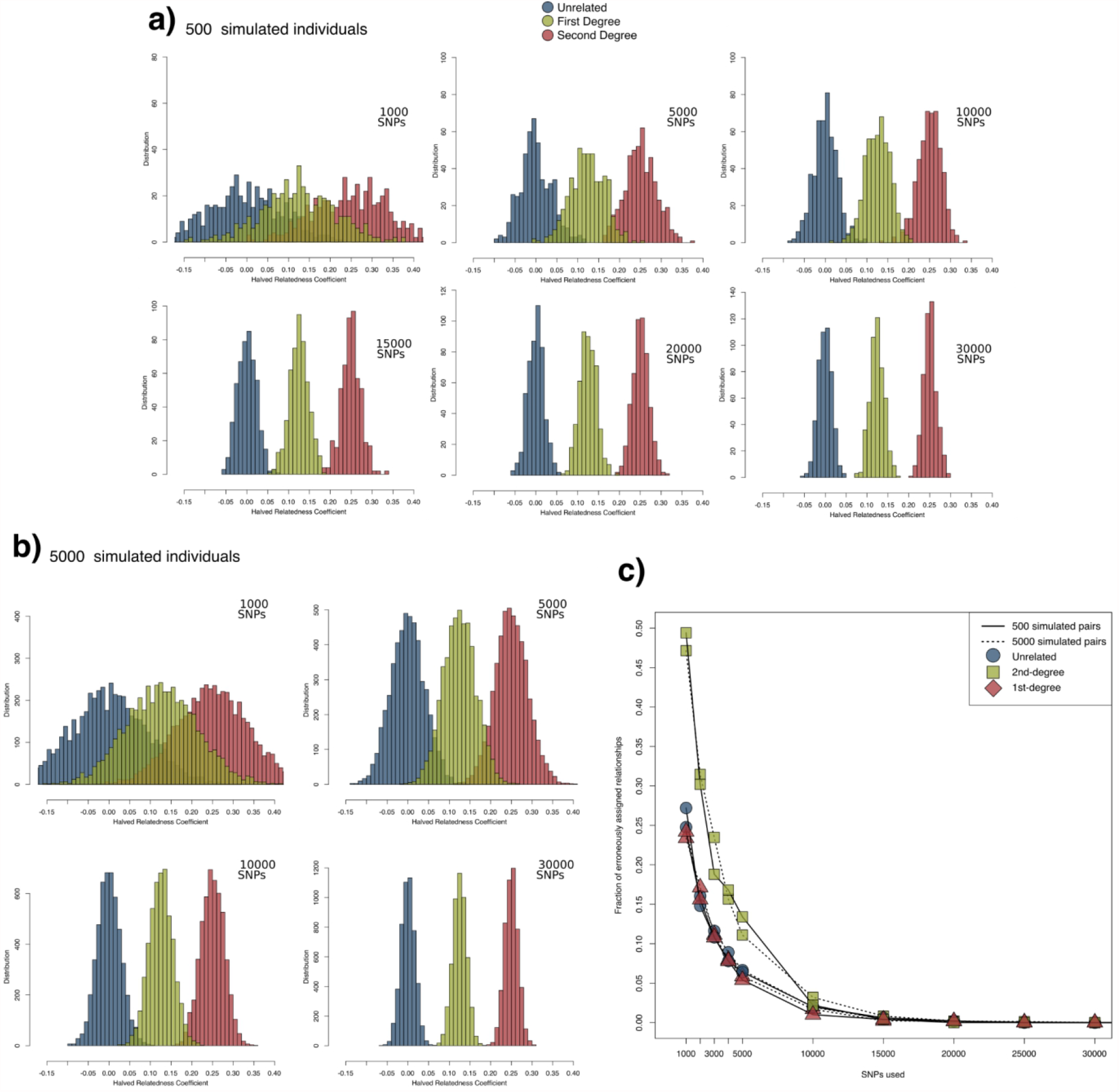
Coefficient distribution ranges for **a)** 500 and **b)** 5000 pairs of simulated individuals using different numbers of SNPs, based on Phase 1 CHS allele frequencies, demonstrating overlaps and the correction of the curves towards the hard thresholds between classes on the higher SNP numbers. **c)** False positive rates, as identified by simulated relationships crossing the thresholds between classes. From 30,000 SNPs no overlap was obtained, even with up to 5,000 simulated pairs of individuals, although higher numbers of simulated pairs would eventually produce an overlap with error rates further tending towards 0.

Increasing the numbers of simulated pairs to induce an expansion of the normal curves and produce overlaps when using large SNP numbers can quickly become very computationally intensive using our pipeline, but our results show that the rates of incorrect estimates based on overlapping ranges are limited to the number of SNPs used and not the number of simulated pairs. In **Figure 3c** we show that these false positive rates stay mostly constant with both 500 and 5000 simulated pairs. By looking at the 2nd-degree curves, which can overlap with both the unrelated and 1st-degree curves, the error rate is on average 47.90% (47.07 to 49.40%, depending on how many pairs were simulated) with 1,000 SNPs, meaning that almost half of the simulated 2nd-degree pairs have their HRC overlapping any of the other classes (**Figure 3a/b, Supplementary Table 3**). However, these rates are reduced to averages of 12.83% on 5,000 SNPs, to 2.81% on 10,000 SNPs, to 0.49% on 15,000 SNPs, and to 0.06% on 20,000 SNPs. For the latter, only 10 out of 5,000 simulated 2nd-degree pairs had a HRC overlapping the other classes’ ranges.

Overall, these simulation results are in agreement with the estimates presented for the CHS population in **Figure 1**, and suggest that using between 15,000 and 25,000 SNPs will likely result in error rates well below 1%. Although different population frequencies are unlikely to produce substantially different error rates on genome-wide shotgun SNP sets, as we report similar error rates using allele frequencies from the 1000 Genomes’ British in England and Scotland (GBR) population (**Supplementary Table 4**), the user is advised to calculate dataset-specific error rates based on simulations using a provided script.

Lastly, the observation in the subsampling experiment of the Koszyce individuals that true 1st-degree relatives were always assigned as 1st-or 2nd-degree relatives based on as few as 4,000 SNPs also finds support in the simulated data. Here, at 5,000 SNPs, the lowest HRC for 1st-degree relatives was 0.095, among all 7100 pairs of simulated individuals, and only 5 times the HRC was below 0.125 (**Supplementary Table 3**).

### Comparison with other relatedness assessment methods used for ancient DNA

We compared two relatedness methods commonly used with low-coverage aDNA data to our method, mainly to evaluate if they were able to produce correct results at similar coverages.

The first method we compared was lcMLkin^5^, which allows the possibility to detect up to 3rd-5th degree relatives^17,18^. For each individual from Koszyce subsampled to the same 1,300,000 and 3,000,000 reads used for testing our method above, we used bcftools to generate genotype likelihoods for ∼22,000,000 variants of the 1000 Genomes Phase 3. However, the number of SNPs overlapping each pair of individuals was always below 300, which is much lower than the minimum suggested amount of 10,000 SNPs in the software’s manual.

The second method we compared was READ, which can detect up to 2nd-degree relatives with coverages as low as 0.1X^3^, although it has also been successfully applied to individuals with 0.03x coverage^15^. Again, we used the same 1,300,000 and 3,000,000 read sets, and, following READ’s manual, we excluded non-polymorphic and low frequency variants by using --maf 0.01 in PLINK^1^.

At the lowest threshold and with default settings, READ produces a “S-like” curve on its graphical output file for the 105 pairwise relationships, which is indicative of different relationships (**Supplementary Figure 3a**). However, all individuals are classified as unrelated and the error bars indicate substantial relatedness class overlaps and high uncertainty. Using user-specified normalisation methods did not help produce a larger spread of estimates, and even with the “max” option only two individuals are classified as 2nd-degree (**Supplementary Figure 3a**). The non-normalised P0 values confirm that there is not a completely clear separation of individuals based on their expected relationship, especially in regards to the unrelated and 2nd-degree classes (**Supplementary Table 5**). Using the 3,000,000 reads threshold there is an apparent better separation but there is still a lack of enough variation to be able to properly identify relatives, with the “max” normalisation setting detecting some 2nd-degree relatives but still unable to detect any 1st-degree relatives (**Supplementary Figure 3b**).

We tried to understand if these results were caused by lack of data or by the use of the genome-wide SNP dataset. To do so, we extracted the Koszyce individuals genotyped to the variants in the 1240K dataset^19^ from the Allen Ancient DNA Resource v44.3 (https://reich.hms.harvard.edu/, accessed on 20.03.2021) and randomly downsampled them to have similar numbers of overlapping SNPs as when we used the 1,300,000 reads and our genome-wide dataset. This corresponded to around 3,300 SNPs per pair. By using SNPs from the 1240K dataset indeed READ is able to correctly assigned all 12 1st-degree relationships, as well as 23 of the 32 2nd-degrees (although here 11 had high uncertainty reflected in values of |Z| < 1) (**Supplementary Figure 3c, Supplementary Table 5**). This suggests that READ can accurately predict a high number of correct relationships with only ∼3300 SNPs from the 1240K dataset, although mostly for the 1st-degree class, and that the use of a curated dataset based on informative SNPs such as this one (as opposed to a genome-wide SNP set) can have a strong influence on the results and the minimum amount of data needed for relatedness estimation. Indeed, in **Supplementary Figure 4** we show that TKGWV2 only requires around 3,000 SNPs to achieve average error rates below 1% when the 1240K dataset is used. However, for samples obtained from genome-wide shotgun data, using the 1240K dataset over a genome-wide dataset to assess kinship is disadvantageous, as the amount of data needed to obtain similarly low error rates (1% with around 3,000 used 1240K SNPs) would necessarily translate into minimum shotgun coverages closer to the more traditional threshold of 0.1X.

## Discussion

We present an updated pipeline for a methodology used to identify the remains of Thomas Kent, an Irish nationalist executed after the Irish Easter Rising, in 1916^4^. The pipeline uses Python, R, and Bash, for a more streamlined and faster process, especially when multiple pairs of individuals are analysed simultaneously. The 105 pairwise relationships between the 15 Koszyce individuals subset to 1,300,000 SNPs ran in 61 minutes, averaging ∼35 seconds per pair, on an Intel Core i7-7700K processor. More importantly, by using genome-wide variants instead of pre-defined SNP sets, this updated pipeline is able to assess kinship relationships with very low error rates using genomic coverages as low as around 0.02x. However, as shown during its application to a set of ancient individuals from Saag et al.^15^, specifically for the pair of individuals X05 -X06, it can also be applied for a genetic relatives pair where one individual has even lower coverage (as low as 0.005x -X06) and the other individual has a substantially higher coverage (for example, 0.03x -X05). In these situations, although the resulting estimates can have some uncertainty due to the small number of common SNPs used, when interpreted together with simulated population sets and false positive/negative rates, they can be a helpful tool in project planning during screening sequencing runs, by informing decisions on whether to further sequence some individuals within a dataset.

With DNA sequencing costs decreasing^20^, aDNA data growing exponentially, and research projects increasingly focusing on analysing the social organisation of populations from specific regions and/or local cemeteries^13,17,18,21–23^, a method for detecting kinship from whole genome shotgun screening data with ultra low coverage is valuable, and can translate into larger numbers of relationships detected. This is true even for individuals that would not be expected to yield sufficient data for population genetic analysis, or for which the required sequencing power would require substantially more funds. Furthermore, as it is common practice to exclude 1st-degree relatives from population genetic analyses that are not focused on local organisation and structure, the detection of such pairs of individuals during initial screening steps can allow researchers to further sequence only the best of the two individuals, and therefore save sequencing power. Our results suggest that from as low as 600,000 aligned reads (0.012x coverage) it is possible to detect 1st-degree relatives with very low false positive rates.

Lastly, by relying on population frequencies calculated from an external population, and requiring no normalisation steps based on unrelated individuals from the population being tested (or genetically similar)^3,7^, TKGWV2 can solve relationships when a single pair of individuals is given as input, or even simultaneously between individuals belonging to different populations and time periods, producing results always within the same expected intervals. We show that, for example, using allele frequency data from modern European populations allowed us to correctly estimate all relationships tested from ancient individuals from that region, potentially allowing the method to be used in any other ancient population where there is substantial DNA coverage from the modern populations they contributed substantial amounts of ancestry to, even if only a single pair of ancient individuals exists. Additionally, we show that this modern population data does not necessarily need to fully match the genetic structure of the ancient individuals being tested, as we produced accurate relatedness estimation results for populations spanning 5,000 years -from Neolithic individuals composed mainly of Early European Farmer and Western Hunter-Gatherer ancestry^4,13,24^, to Bronze Age and Medieval individuals with substantial amounts of Steppe-related ancestry^e.g. 14,15^.

An overall caveat of using modern population frequencies on ancient individuals, however, is that, when applying this approach to any previously untested ancient population, a confirmation analysis on known ancient related individuals needs to be performed using frequencies from the closest modern populations available. This can be challenging for regions or periods for which there is a lack of published ancient relatives. Nevertheless, with the ongoing exponential increase in availability of both modern and aDNA data around the world, our method has the potential to be applied to the great majority of situations encountered by palaeogeneticists.

## Acknowledgements

We would like to thank Kendra Sirak for comments and suggestions on this manuscript; Torsten Günther for support in using READ and interpreting its below-threshold results; and Alissa Mittnik for discussions before submission.

## Author Contributions

D.F., P.G. and R.P. conceptualized the manuscript. O.C. analysed the data to produce Supplementary Figure 1. D.F. developed the original method’s update, wrote the bioinformatics package, analysed the data, performed simulations, and wrote the manuscript with input from all co-authors.

## Data Availability

All genomic data used in this manuscript have been previously published. The modern data from the 1000 Genomes was downloaded from ftp://ftp.1000genomes.ebi.ac.uk, and the ancient data was downloaded from https://www.ebi.ac.uk/, using the accession numbers found in the original publications.

## Bibliography

1. Chang, C. C. et al.. Second-generation PLINK: rising to the challenge of larger and richer datasets. Gigascience 4, 7 (2015).

2. Manichaikul, A. et al.. Robust relationship inference in genome-wide association studies. Bioinformatics 26, 2867–2873 (2010).

3. Monroy Kuhn, J. M., Jakobsson, M. & Günther, T. Estimating genetic kin relationships in prehistoric populations. PLoS One 13, e0195491 (2018).

4. Fernandes, D. et al.. The Identification of a 1916 Irish Rebel: New Approach for Estimating Relatedness From Low Coverage Homozygous Genomes. Sci. Rep. 7, 41529 (2017).

5. Lipatov, M., Sanjeev, K., Patro, R. & Veeramah, K. Maximum Likelihood Estimation of Biological Relatedness from Low Coverage Sequencing Data. bioRxiv (2015) doi:10.1101/023374.

6. Korneliussen, T. S. & Moltke, I. NgsRelate: a software tool for estimating pairwise relatedness from next-generation sequencing data. Bioinformatics 31, 4009–4011 (2015).

7. Olalde, I. et al.. The Beaker phenomenon and the genomic transformation of northwest Europe. Nature 555, 190–196 (2018).

8. Wang, J. Effects of sampling close relatives on some elementary population genetics analyses. Molecular Ecology Resources 18, 41–54 (2018).

9. Olalde, I. & Posth, C. Latest trends in archaeogenetic research of west Eurasians. Curr. Opin. Genet. Dev. 62, 36–43 (2020).

10. 1000 Genomes Project Consortium et al. A global reference for human genetic variation. Nature 526, 68–74 (2015).

11. Queller, D. C. & Goodnight, K. F. ESTIMATING RELATEDNESS USING GENETIC MARKERS. Evolution 43, 258–275 (1989).

12. Schlauch, D. Methods for Estimating Hidden Structure and Network Transitions in Genomics. (Harvard University, 2017).

13. Schroeder, H. et al.. Unraveling ancestry, kinship, and violence in a Late Neolithic mass grave. Proc. Natl. Acad. Sci. U. S. A. 116, 10705–10710 (2019).

14. Margaryan, A. et al.. Population genomics of the Viking world. Nature 585, 390–396 (2020).

15. Saag, L. et al.. The Arrival of Siberian Ancestry Connecting the Eastern Baltic to Uralic Speakers further East. Curr. Biol. 29, 1701–1711.e16 (2019).

16. Hardy, O. J. Estimation of pairwise relatedness between individuals and characterization of isolation-by-distance processes using dominant genetic markers. Mol. Ecol. 12, 1577–1588 (2003).

17. Mittnik, A. et al.. Kinship-based social inequality in Bronze Age Europe. Science 366, 731–734 (2019).

18. Cassidy, L. M. et al.. A dynastic elite in monumental Neolithic society. Nature 582, 384–388 (2020).

19. Mathieson, I. et al.. Genome-wide patterns of selection in 230 ancient Eurasians. Nature 528, 499–503 (2015).

20. Wetterstrand, K. DNA Sequencing Costs: Data. National Human Genome Research Institute https://www.genome.gov/about-genomics/fact-sheets/DNA-Sequencing-Costs-Data (2021)

21. Fernandes, D. M. et al.. A genetic history of the pre-contact Caribbean. Nature 590, 103–110 (2020).

22. Amorim, C. E. G. et al.. Understanding 6th-century barbarian social organization and migration through paleogenomics. Nat. Commun. 9, 3547 (2018).

23. Sirak, K. A. et al.. Social stratification without genetic differentiation at the site of Kulubnarti in Christian Period Nubia. bioRxiv (2021) doi:10.1101/2021.02.17.431423.

24. Brace, S. et al.. Ancient genomes indicate population replacement in Early Neolithic Britain. Nat Ecol Evol 3, 765–771 (2019).

