## Supplementary Information for "TKGWV2: An ancient DNA relatedness pipeline for ultra-low coverage whole genome shotgun data"

Includes:

**Supplementary Figure 1**

Possible pedigree explaining the relationships between individuals X13 (0.0147x), X14 (0.307), and V16 (0.220). Bold lines represent newly described relationships.

**Supplementary Figure 2**

Simulated distribution ranges for 7 pairs of individuals from Saag et al. 2019, with less than 10,000 common SNPs. HRC shown as yellow lines, and posterior probabilities for each of the three curves/classes as white text.

**Supplementary Figure 3**

Pairwise relationships for the Koszyce individuals subsampled to 3 different numbers of reads or common SNPs, using the “max” (plus “median” in a)) normalization setting in READ. a) Using 1,300,000 aligned reads per individual and genome-wide SNPs, using our method. Coloured points and right scale correspond to READ results using the default normalization setting (“median”). b) Same as a) but using 3,000,000 aligned reads, and showing only “max”. c) Results using an average of 3,300 SNPs from the 1240K dataset per pair.

**Supplementary Figure 4**

False positive rates, as identified by simulated relationships crossing the thresholds between classes when using the 1240K SNP set. From around 3,000 SNPs used error rates are below 1%.

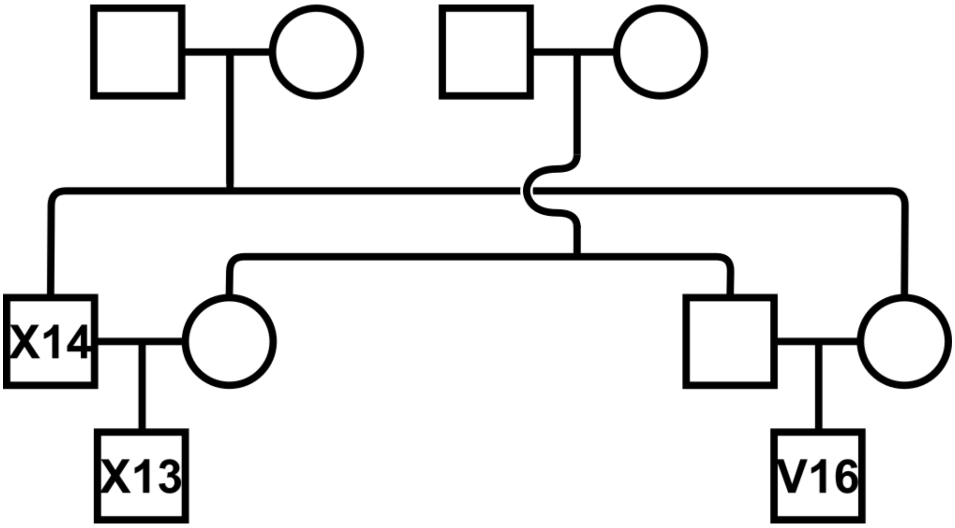


**Supplementary Figure 1.** Possible pedigree explaining the relationships between individuals X13 (0.0147x), X14 (0.307), and V16 (0.220). Bold lines represent newly described relationships.

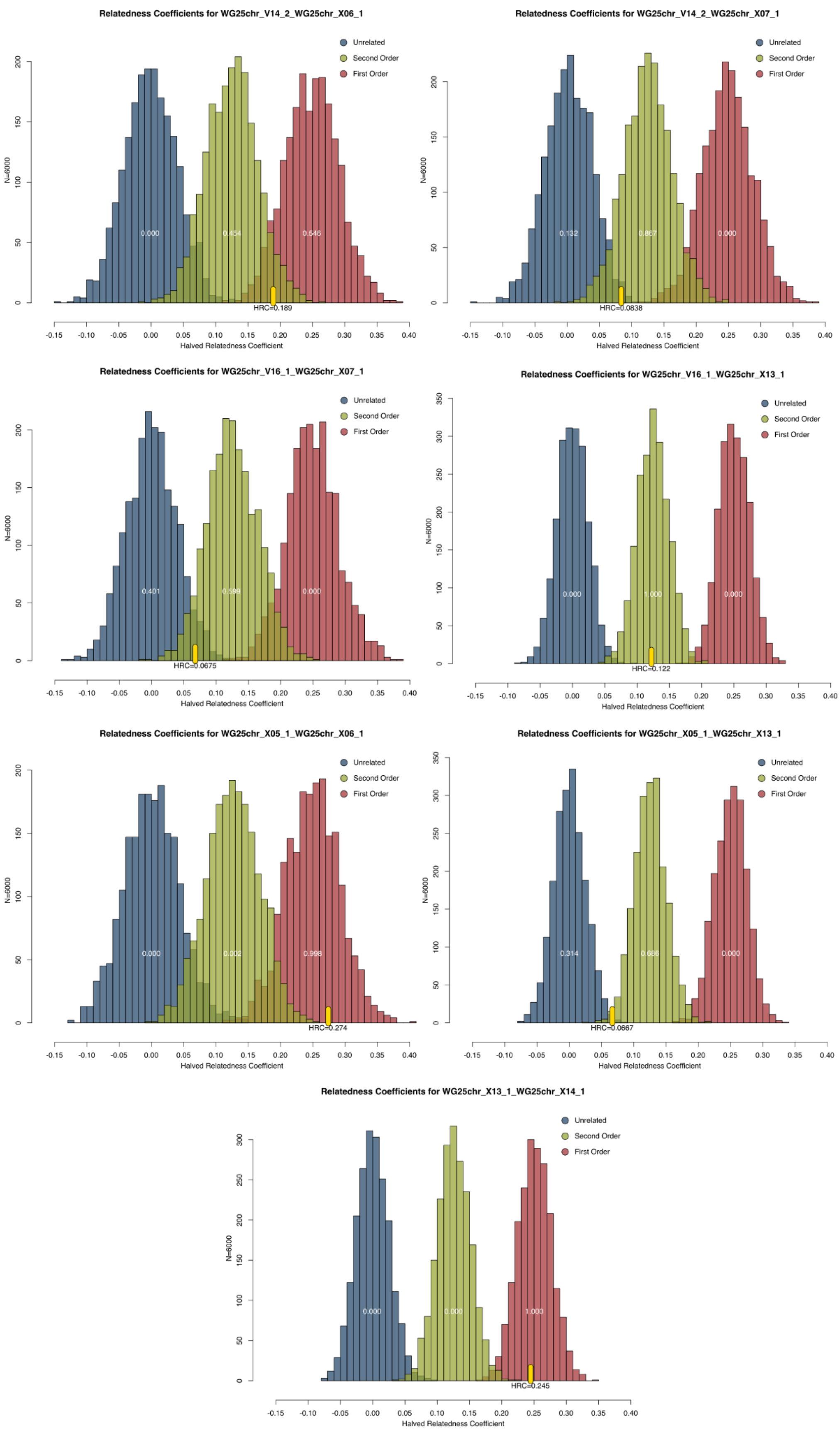


**Supplementary Figure 2.** Simulated distribution ranges for 7 pairs of individuals from Saag et al. 2019, with less than 10,000 common SNPs. HRC shown as yellow lines, and posterior probabilities for each of the three curves/classes as white text.

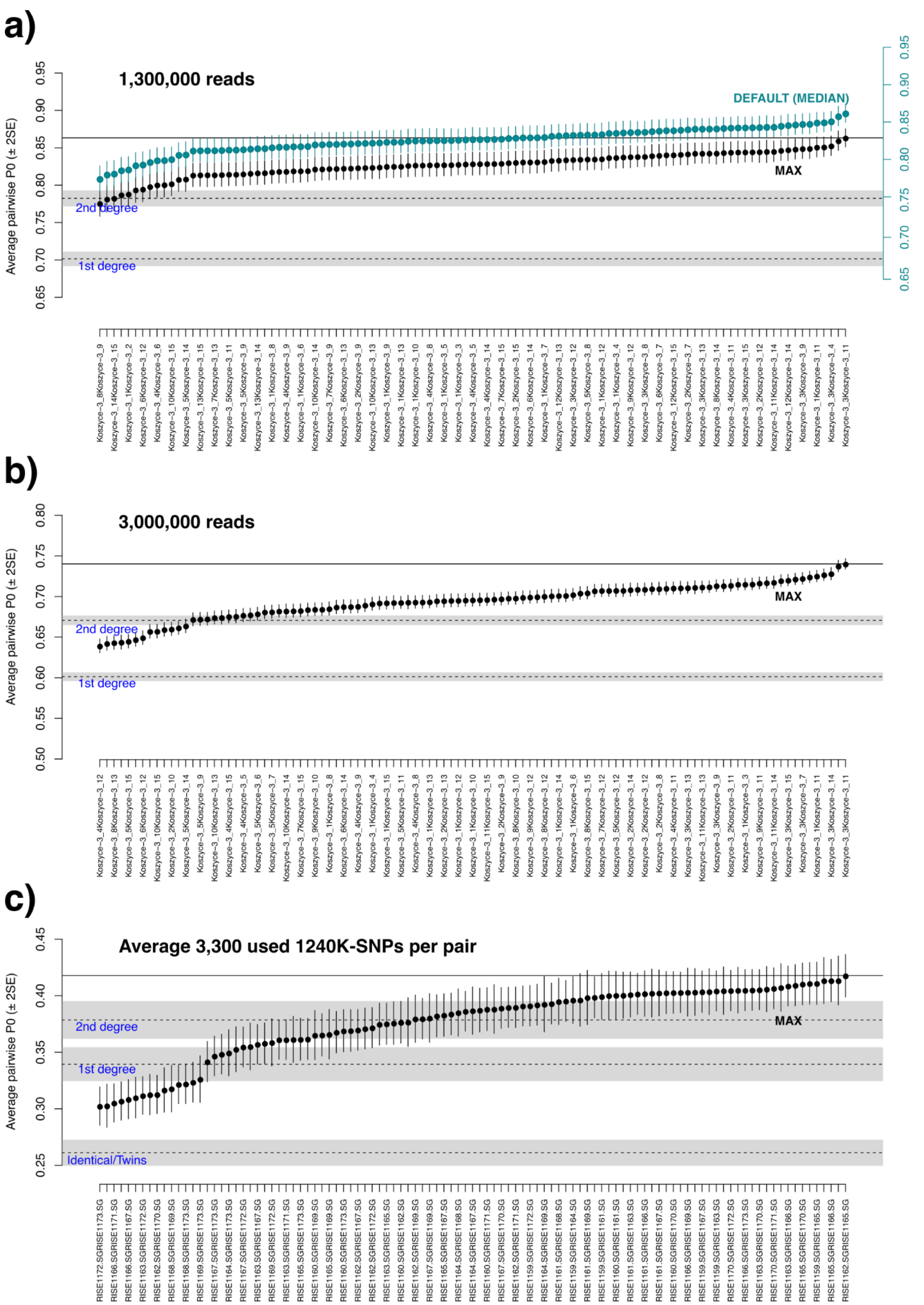


**Supplementary Figure 3.** Pairwise relationships for the Koszyce individuals subsampled to 3 different numbers of reads or common SNPs, using the "max" (plus "median" in **a)**) normalization setting in READ. **a)** Using 1,300,000 aligned reads per individual and genome-wide SNPs, using our method. Coloured points and right scale correspond to READ results using the default normalization setting ("median"). **b)** Same as **a)** but using 3,000,000 aligned reads, and showing only "max". **c)** Results using an average of 3,300 SNPs from the 1240K dataset per pair.

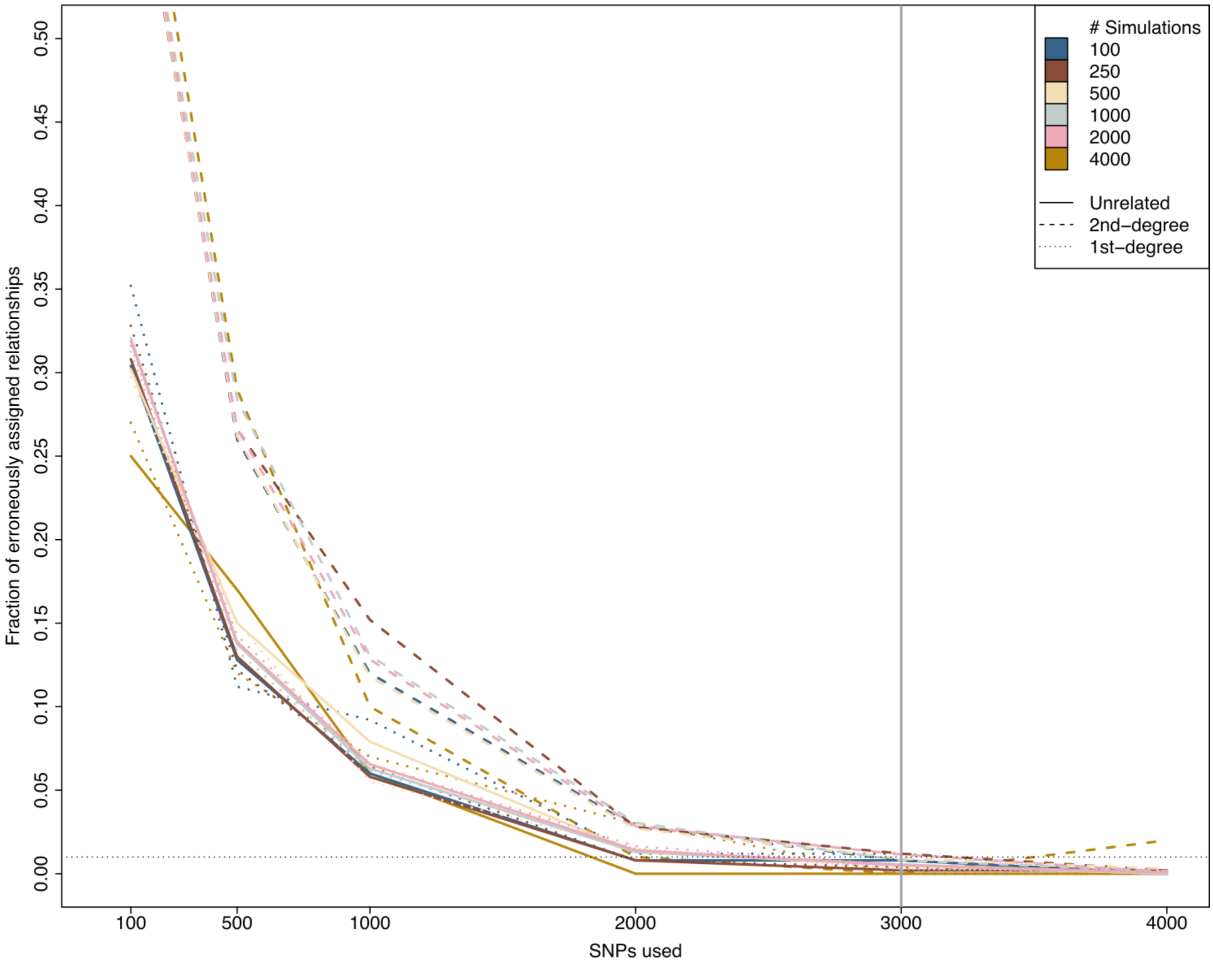


**Supplementary Figure 4.** False positive rates, as identified by simulated relationships crossing the thresholds between classes when using the 1240K SNP set. From around 3,000 SNPs used error rates are below 1%.
